# Robust bacterial co-occurence community structures are independent of *r*- and *K*-selection history

**DOI:** 10.1101/2021.08.04.455033

**Authors:** Jakob Peder Pettersen, Madeleine S. Gundersen, Eivind Almaas

## Abstract

Selection for bacteria which are *K*-strategists instead of *r*-strategists has been shown to improve fish health and survival in aquaculture. We considered an experiment where microcosms were inoculated with natural seawater and the selection regime was switched from *K*-selection (by continuous feeding) to *r*-selection (by pulse feeding) and vice versa. We found the networks of significant co-occurrences to contain clusters of taxonomically related bacteria having positive associations. Comparing this with the time dynamics, we found that the clusters most likely were results of similar niche preferences of the involved bacteria. In particular, the distinction between *r*- or *K*-strategists was evident. Each selection regime seemed to give rise to a specific pattern, to which the community converges regardless of its prehistory. Furthermore, the results proved robust to parameter choices in the analysis, such as the filtering threshold, level of random noise, replacing absolute abundances with relative abundances, and the choice of similarity measure. Even though our data and approaches cannot directly predict ecological interactions, our approach provides insights on how the selection regime affects the composition and workings of the microbial community, providing a basis for aquaculture experiments targeted at eliminating opportunistic fish pathogens.

## Introduction

In aquaculture, the fish is in close contact with its environmental microbiome^1^. Fish larvae are at an especially vulnerable life stage, and the larvae’s high death rate cause economic problems in the aquaculture industry^2,3^. Research during the last decades has uncovered that the bacterial composition of the larvas’ environment affects larval survival, and both detrimental and favourable host-microbe interactions have been identified. This strongly suggests that the environmental microbiome of larvae can be manipulated to improve health and survival^4,5^. Since detrimental interactions are linked to opportunistic pathogens that outcompete favourable mutualistic bacteria and cause disease when in excess^4,5^, a viable approach for microbiome control is to select against the opportunistic pathogens by instead selecting for more favourable bacteria. The concept of *r*- and *K*-strategists, introduced in microbial ecology by Andrews and Harris^6^, provides a guideline for how to implement such a selection. Most opportunists are *r*-strategists, meaning that they grow rapidly when resources are in surplus. On the other hand, *r*-strategists complete poorly when the environment is resource-limited. In such a competitive environment, the slow-growing *K*-strategists will quickly dominate due to their high substrate acquiring rate and high yield^1,7^.

To illustrate the concept of *r/K*-selection, we consider an example from recirculating aquaculture systems (RAS): UV light or ozone has been frequently applied in RAS to disinfect the rearing water. The idea behind this procedure is to remove the fish pathogens such that the fish will stay healthier^8,9^. Counter-intuitively, experiments have shown that this disinfection actually led to poorer fish health^10,11^. Why? As earlier mentioned, microbes can have both favourable and detrimental interactions with the fish. These interactions are not related to the bacterial load, but rather the composition. UV light reduces the bacterial load and kills potential pathogens and protective bacteria alike. Still, some bacteria will remain in the system and will now have far fewer competitors for the same amount of resources. In turn, the opportunistic *r*-strategists will quickly proliferate on this feast and subsequently cause disease to the fish, while the *K*-strategists will have too low maximum growth rates to compete^7^.

This example teaches us that it is important to understand life strategies of bacteria and how they influence community composition and dynamics. Getting a deeper understanding of these concepts is likely to resolve problems related to fish microbiota in aquaculture and other microbe-dominated processes.

In this article, we aimed at unveiling the structure and community dynamics in *r*- and *K*-selected communities using similarity measures and network inference. We hypothesized that such a tool could separate bacteria based on their growth preferences and, hence, be useful to characterise and identify the *r*- and *K*-strategists. In particular, we were interested in answering whether positive or negative associations were the most dominant, and how they contributed to the overall network structure.

To study interactions in *r*- and *K*-selected communities, we analysed a 2×2 factorial crossover microcosm experiment that tested varying feeding regime and resource availability (high/low).^12^ Briefly, half of the microcosms were pulse-fed resources which promotes the growth of *r*-strategists, whereas the other half received a steady, continuous supply of nutrients promoting *K*-strategists. We hereafter refer to these selection regimes as *r*- and *K*-selection. The bacterial communities were sampled and characterised through 16S ribosomal DNA (rDNA) sequencing^13–15^ at 18 time-points over a 50 day period. In contrast to many other microbiome experiments, in this experiment the authors determined bacterial density by flow cytometry^12^, even though these data were not mentioned in the article. As sequencing on its own only provides relative abundances of microbes in each, knowing bacterial densities allowed us to estimate absolute abundances.

What made this experiment particularly suited for our analysis, was that approximately halfway into the experiment (i.e. between day 28 and 29), the *r*- and *K*-selection regimes were switched such that each microcosm was subjected to both selection regimes. This allows for some unique network conclusions to be drawn as we may, with much higher certainty, relate our network results to the *r*- and *K*-selection. Particularly, we were interested in investigating whether the *r*- and *K*-selection resulted in reproducible results and whether such *r*- and *K*-selection gave the the communities memories and resilience. For microbial cultivators who want to control the microbiome, understanding the effect this *r/K*-selection regime-switch is important.

## Results

To investigate the bacterial community structure and dynamics in *r*- and *K*-selected communities, we assessed the co-occurrence patterns of 1,537 operational taxonomic units^16^ (OTUs) observed in the microcosm experiment. This 16S-rDNA dataset consisted of 202 samples from 12 microcosms cultivated over 50 days. Note that, 6 of the microcosm were *r*-selected, and 6 were *K*-selected. Between day 28 and 29, the *r/K*-selection regime was switched such that *r*-selected communities now were *K*-selected (the RK-group) and vice versa (the KR group). Furthermore, the microcosms varied by the amount of resources supplied, high (H) and low (L). However, exploratory analysis of the dataset did not indicate any relevant effect of the resource supply, and hereafter we will only focus on the *r*- and *K*-selection regimes.

### Similarity measurements and network inference

We assessed the co-occurrence patters between the OTUs using seven similarity measurements and varying levels of random noise, OTU filtering, and type of abundance (relative/absolute). Here, we present the results for the Spearman correlation measure with a low level of random noise, low OTU filtering threshold and absolute abundances (see the Methods’ section for more details). We decided to focus on the rank-based Spearman correlation because it is widely applied for detecting associations^15,17,18^. We will later discuss the robustness of these results by contrasting and comparing with other similarity measures and parameter choices.

We wanted to create a network of the pairwise associations between the OTUs and thus had to determine which edges to include. Selecting a hard threshold for the *q*-value for an interaction to be statistically significant (for instance at *q* ≤ 0.05), is not an easy choice^19^. We, therefore, illustrate the number of significant interactions over a range of threshold *q*-values up to 0.05 (Fig.1). From this figure, we see that there is no obvious cutoff.

**Figure 1.**
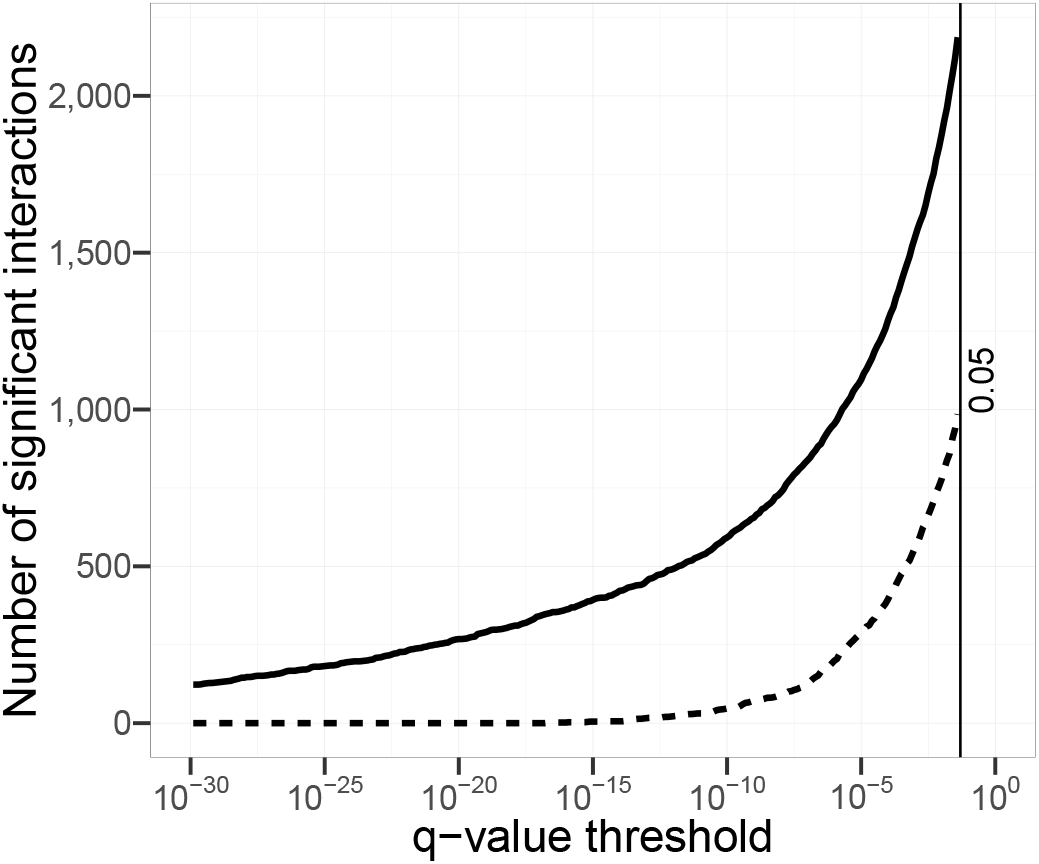
The cumulative number of significant interactions as a function of the critical *q*-value threshold considered. The solid line signifies positive interactions detected, while the dashed line represents the number of negative interactions.

There were in total 3,250 interactions having *q* ≤ 0.05, of which 1,679 were 0.05 ≥ *q* ≥ 10^−4^ and 639 with *q* < 10^−10^. Therefore, we determined 500 edges to be a reasonable balance between selecting high-significance edges and a network with lower average node connections. With this setting, the effective *q*-value hreshold became 5.0 .10^−13^. The resulting network modules were labelled using the walktrap algorithm with 20 steps^20^ (Fig.2).

**Figure 2.**
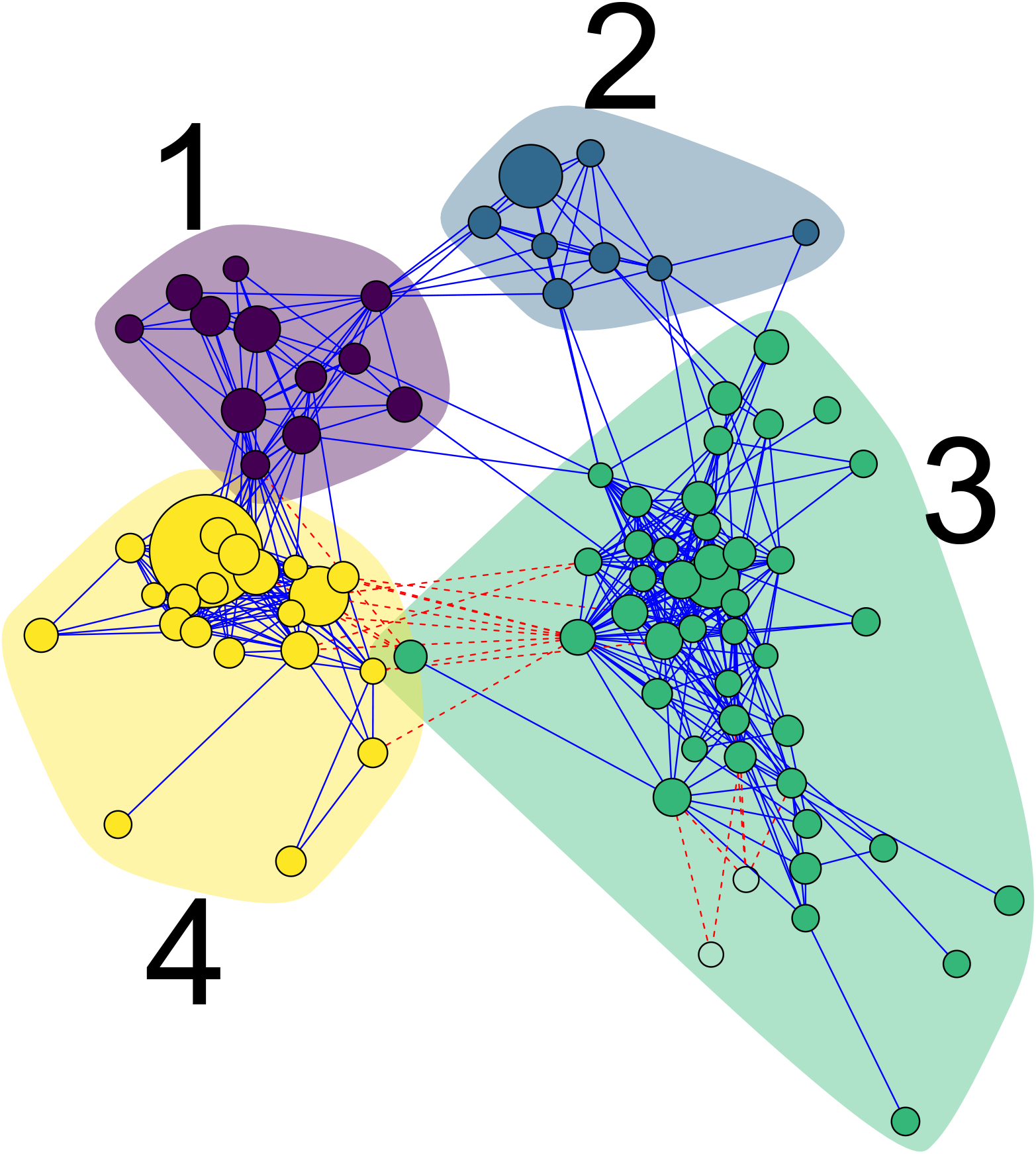
Module-labelled network of the 500 most significant interactions in the *r/K*-selection-switch dataset. Each of the 86 nodes is an OTU, while each edge corresponds to a statistically significant association between the OTUs. Blue solid edges indicate positive interactions, whereas red dashed edges indicate negative interactions. The node-sizes are scaled logarithmically according to overall mean abundance.

### Phylogenetic clustering within the modules

The co-occurance network analysis clustered the OTUs into four distinct modules (Fig.2). Also, two OTUs were not assigned any module (shown as colourless nodes inside module 3) as they were connected to the rest of the network with negative interactions only. Module 3 and 4 stood out as the most interesting modules for two reasons. First, they had the largest number of nodes. Second, there were negative links between the modules, suggesting antagonistic relationship between the modules. For these two main modules we observed a large number of positive links within the modules, but negative edges between them. Therefore we wondered whether the modules were phylogenetically clustered. Indeed, we observed a clear pattern between OTU module membership and phylogenetic classification (Fig.3 and Supplementary Table S1). Module 3 consisted primarily of *Alphaproteobacteria* (including *Rhodobacteraceae*) and *Flavobacteria*, whereas module 4 was dominated by *Gammaproteobacteria* (including *Colwellia* and *Vibrio*). Hence, the modules were enriched in phylogenetically related bacteria.

**Figure 3.**
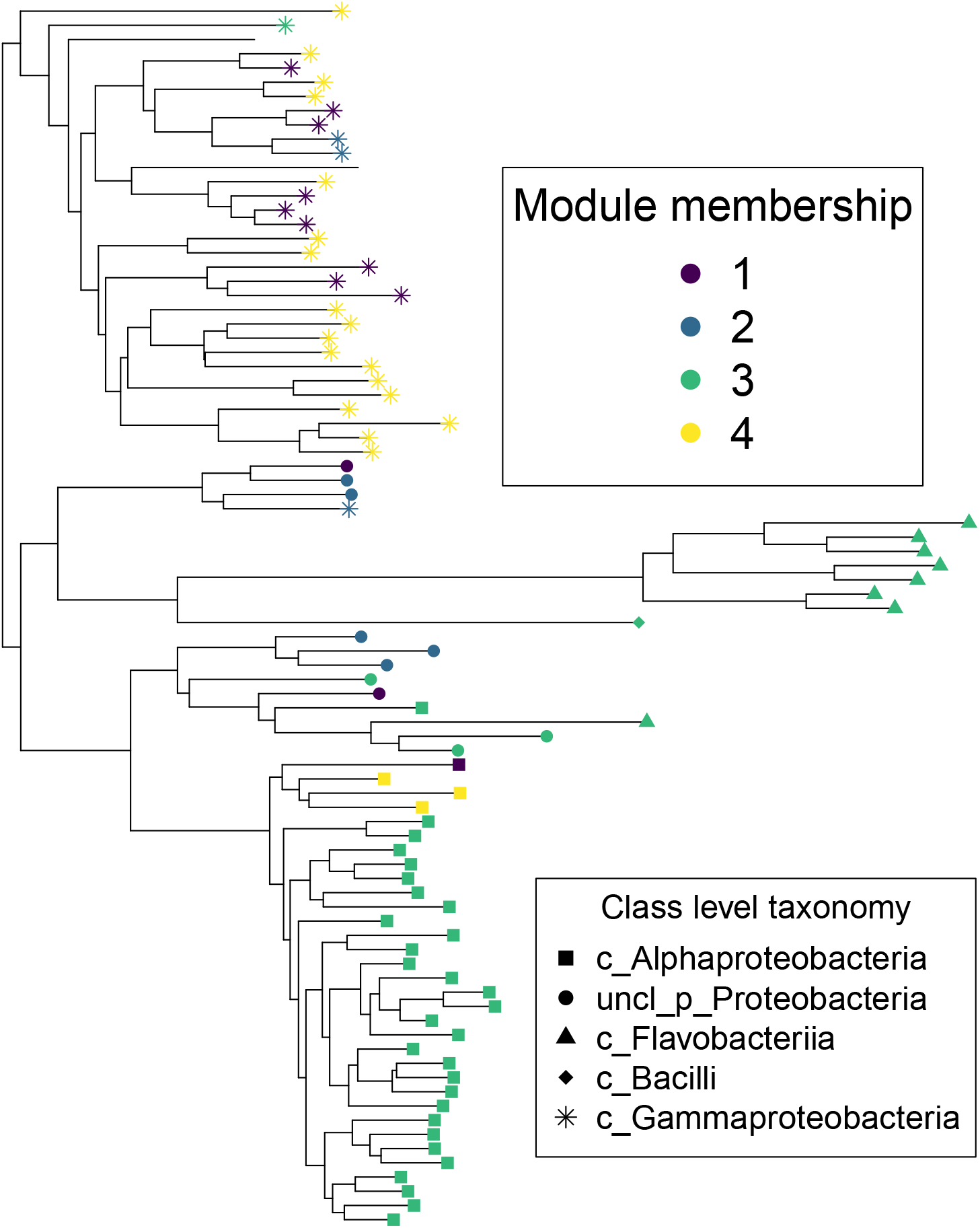
The phylogentic tree of the 86 OTUs from Fig.2. together with the class level taxonomical assignment. Point colour indicates module membership, whereas the shape indicates class level taxonomical assignment. Notice that there are some inconsistencies between the phylogenetic tree and the assigned taxonomy.

### Temporal trajectories of the microbial communities

After having observed the network modules, we were interested in understanding the co-occurrence structures and how it influenced community dynamics. To investigate the community dynamics within the microbial communities, we plotted the Bray-Curtis PCoA-ordinations of the samples and observed the successional trajectories of each microcosm (Fig.4).

**Figure 4.**
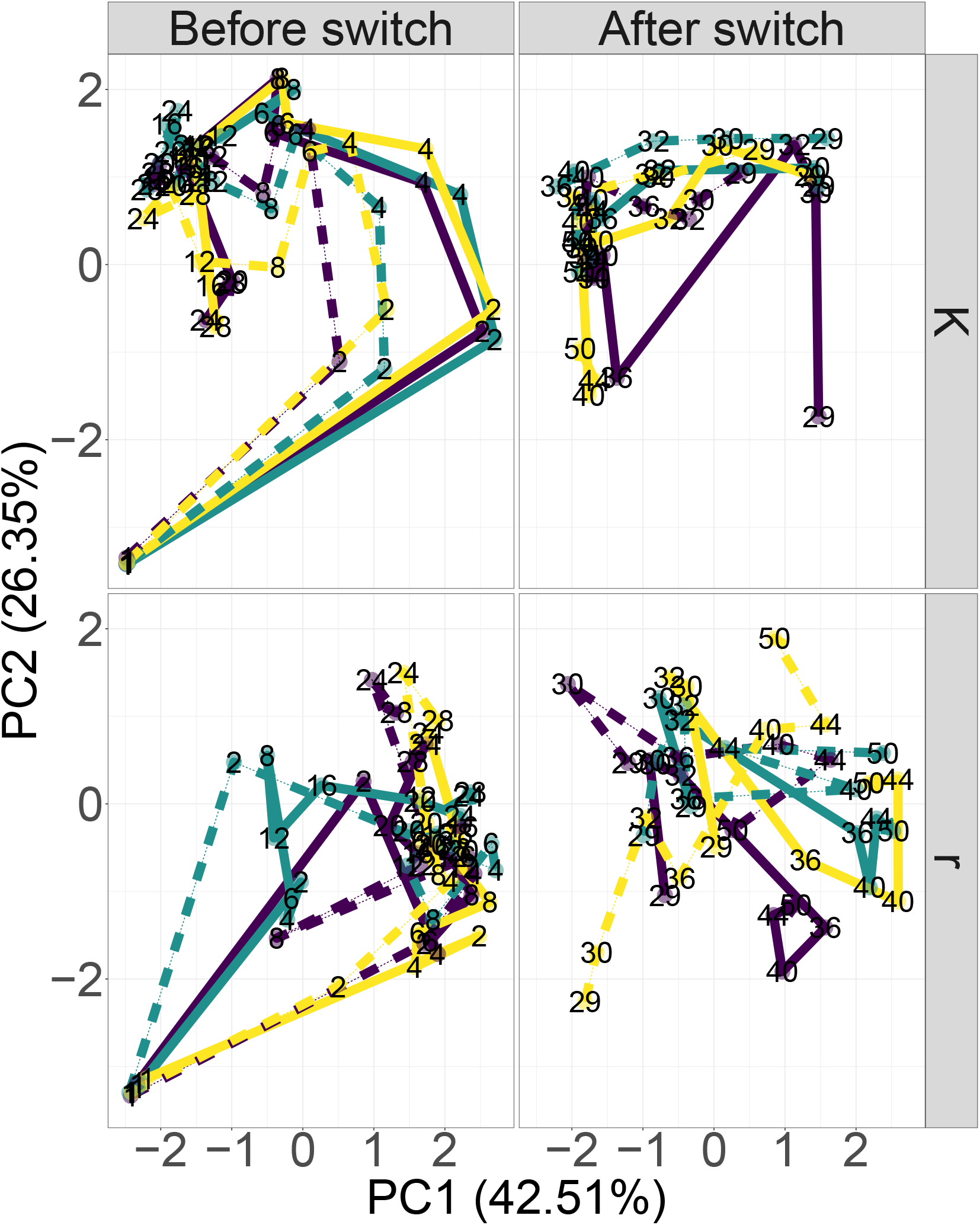
PCoA ordination of Bray-Curtis distances between samples showing the time trajectories for each microcosm. The single ordination was faceted vertically based on the state of selection regime at the time of sampling (being *r* or *K*-selection), and horizontally to highlight temporal trends. Solid and dotted lines indicate high (H) and low (L) nutrient supply, respectively. The labels indicate the day of sampling, whereas the line colours are purely to visually distinguish the replicate time series.

From this time trajectory plot, we observed that microcosms undergoing *K*-section converged towards the upper-left area in the plot, whereas microcosms under *r*-section converged the middle-right area. This effect seemed independent of the experimental period and of pre-existing experimental conditions. Consequently, in this respect the communities did not seem to have any memory-effect that gave rise to resistance against changes in composition. As the *r/K*-selection regimes resulted in clustered communities, we aimed at investigating how the network arose from these dynamics.

### *r/K*-strategist network patterns

We further investigated what influenced the dynamics of the community, and conversely the dynamics’ contributions to the overall network network in Fig.2. For this, we visualised the rank-based *z*-scores for selected days during the experiment for high nutrient supply (Fig.5). For low nutrient supply, the results were very similar and is thus not discussed any further (see Supplementary Fig. S1 for further details).

**Figure 5.**
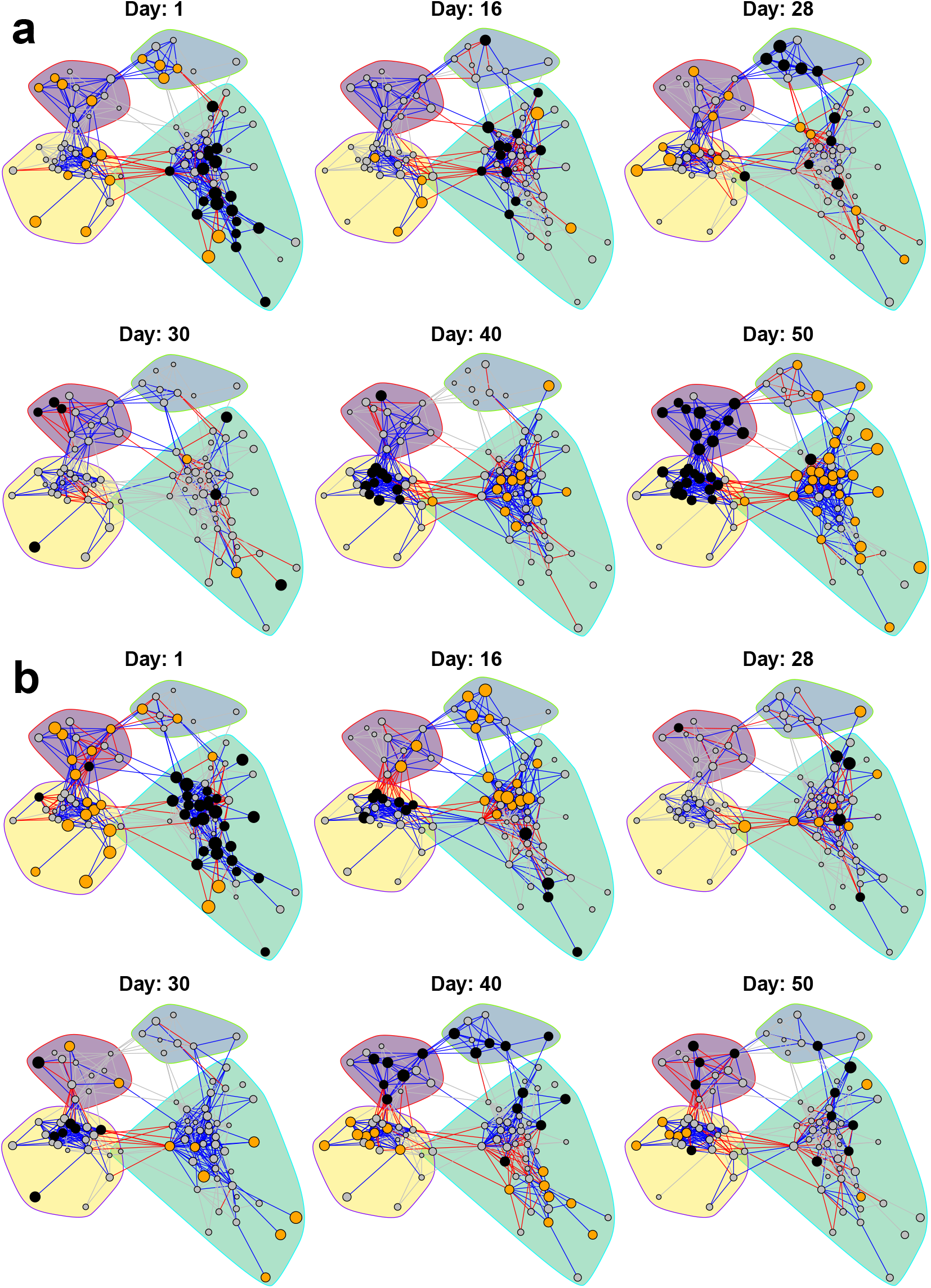
Dynamic visualisation of the network in Fig.2, for **a)** the RK selection group and **b)** the KR selection group for high (H) nutrient supply. Nodes are coloured according to the corresponding OTUs’ abundance compared to its overall mean for all sampling days, represented by its *z*-score. Orange, grey and black nodes mean higher, about the same or lower abundance than its mean, respectively. The edges are coloured by the product of the nodes’ *z*-scores. This means that blue and red edges contribute to positive and negative association across the time series, respectively. The grey edges indicate that no major contribution to neither positive nor negative association is made. As we want to emphasize the orange and black nodes, the nodes with higher absolute *z*-scores are larger.

There were some obvious patterns that were apparent when investigating the temporal networks, especially with regards to module 3 and 4. The abundance of the OTUs in module 3 increased during *K*-selection, while the ones in module 4 had the opposite trend and had high abundances during *r*-selection. Hence, within each module, the OTUs had coordinated abundance patterns leading to positive inferred interactions. On the other hand, between module 3 and 4, the abundance patterns were anti-coordinated such that we obtained negative interactions.

We expected the dataset to display two time periods of instability: The first at the start of the experiment when the microbial community would adapt to lab-culture conditions, and the second disturbance instability after switching the selection regime, after day 28. During these unsteady periods, we expected more instability and less coordination between OTUs belonging to the same module. This in turn, would contribute to negative interactions or weaken the positive ones. However, this expectation was only partially fulfilled because we observed negative edges within modules in Fig.5 also outside the two predicted periods of instability. One potential factor contributing to instability at the beginning of the experiment, was the fact that the oligotrophic seawater was introduced to high amount of nutrients, favouring *r*-strategists to proliferate even if the feeding was continuous.

### Network robustness

If our results were different when changing parameters, the conclusions would be less likely to give us any real insight into how the communities actually behave. Therefore, we checked the robustness of the chosen parameters, changing one at a time while keeping all other parameters constant. Increasing the levels of random noise from low to medium (see Methods section) did not give any substantial difference in terms of significant interactions. Some cosmetic changes were visible due to different color labeling of communities and orientations of plots (details in Supplementary Section S2). Exchanging estimated absolute abundances with relative ones gave higher proportion of negative interactions and different assignments of OTUs into modules (see Supplementary Section S3). However, the greater trends in the results stay the same, such as clustering based on phylogeny and the considerable change in the community behaviour after switching selection regime.

Selecting a more stringent OTU filtering cutoff only has a minor consequence on the results, at the level of cosmetic changes in the plots (see Supplementary Section S4 for details). On the other hand, we notice a more pronounced effect when replacing the Spearman correlation by Pearson correlation. This is not surprising, since Spearman is non-parametric and Pearson measures degree of linear co-occurrence. In this case (Pearson), we got far fewer negative significant interactions for the same *q*-value, none of which are among the 500 most significant ones. Still, modules of phylogenetically related OTUs are present, and the selection regime still seems to explain the modules (Supplementary Section S5).

## Discussion

In literature, challenges of microbial datasets such as sparsity, compositionality and habitat filtering have been addressed and solutions proposed for finding ecological interactions^21–25^. Despite the fact that predictions from ecological interaction-inference tools have been successfully validated in some cases^26,27^, any universally accepted gold standard of finding ecological microbial interactions is not yet agreed upon. Furthermore, some reviews assessing existing methods for inferring ecological interactions have demonstrated that current methods have far too low predictive power, and more refined approaches specifically designed to cope with difficulties in microbial datasets have failed to perform better than the basic ones^18,28^. Hence, we believe that our choice of using the relatively simple ReBoot^23^ procedure is reasonable, even though the approach in and of itself is somewhat coarse-grained.

We observed that our correlation networks clustered the OTUs according to taxonomy and niche preferences as a result of selection for *K*- and *r*-strategists. The finding that taxonomy and niche preferences dominate co-occurrence patterns is in line with work by Chaffron *et al.^29^* who produced similar results from samples stored in a ribosomal RNA database. Along the same line, Bock et. al^30^ also noted that many of the interactions in a correlation network occur between closely related species when studying bacterial and protist communities in European lakes. Bacteria with similar niches are expected to be competitors and, hence, have negative interactions with each other. However, the effect of habitat filtering will create positive correlations between species with similar niches that are often stronger than those arising from ecological competition^31–33^. The same reasoning goes for taxonomical relatedness, as closely related organisms often belong to the same niche and have similar functions. This favours positive interactions within modules, whereas we, to a lesser extent got negative interactions between modules where the growth requirements are different.

Moreover, we have not undertaken any attempt to deal with indirect interactions. This means that two OTUs can appear with a strong (correlation) link even though they have no direct effect on each other, but instead interact with a third OTU. Consequently, it is challenging to determine causality when working with inferred interactions. Also, such indirect effects can be caused by environmental variables and biological entities not taken into account, such as protists and bacteriophages.

For reasons mentioned above, our results are not meant to directly represent real ecological interactions. Nevertheless, our results are interesting from a fish-health perspective, as they show that selection regime can control community composition. In terms of *r*- and *K*-selection, literature consider the orders *Alteromonadales* and *Vibrionales* represented in module 4 as *r*-strategists, whereas the *Rhodobacteraceae* in module 3 are considered *K*-strategists^34^. Additionally, *Vibrio* strains are known to cause disease in fish, whereas *Rhodobacteraceae* bacteria have been shown to protect against *Vibrio* infections through competition^35–38^. This agrees with and extends prior knowledge that *K*-selection is a potent tools for improving fish health and survival^1,7^.

The long-term behaviour of the community did not appear to depend on its prehistory. Potentially, this means that changing the microbiota from a detrimental to a healthy state in a running aquaculture facility requires the same measures as ensuring a healthy microbiota for a new facility. Furthermore, the trustworthiness of the results is strengthened by their robustness to changes in parameter settings, such as filtering cut-off, amount of random noise, type of abundance, and similarity measure.

This experiment was conducted in an artificial setting without any fish of which the health could be tracked. Furthermore, we do not know whether up-scaling and broader exposure could change the workings of the microbial community. Therefore, follow-up studies could be implemented in realistic aquaculture settings, perhaps such as a RAS facility, to investigate whether switching between *K*-selection and *r*-selection will yield the same community dynamics as described in this paper. Additionally, such an experiment would provide opportunity to investigate possible connections between the state of the microbial community and the health of the fish.

We acknowledge that there exist alternative approaches one could follow. For instance, treating the OTUs as discrete units is a bit misleading. As the results show, closely related OTUs often occur together, so it could make more sense to treat the bacteria as a taxonomical continuum. A novel approach based on amplicon sequence variants (ASVs) avoids the clustering of OTUs altogether by considering each individual unique read as an own entity^39^. The phylogenetic relatedness between ASVs could then be used as a constraint for finding co-occurrence patterns. In addition, incorporating environmental information, such as organic nutrient load, salinity, and temperature, would be useful because this allows us to better predict how the desired *K*-selection should be obtained. Joint Species Distribution Models (JSDMs)^40,41^ might have this useful potential to account for both species interaction, environmental factors, and taxonomical relatedness. However, its use in microbial ecology is still in its early stages and time dynamics are not yet embedded into the framework^42,43^.

## Methods

### Quantification of bacterial density

For each sample, the bacterial density was quantified using flow cytometry (BC Accuri C6)^12^. In brief, the bacterial communities were diluted in 0.1× TE buffer, mixed with 2× SYBR Green II RNA gel stain (ThermoFisher Scientific) and incubated in the dark at room temperature for 15 minutes. Then, each sample was measured for 2.5 minutes at 35 μLmin^−1^ with an FL1-H (533/30 nm) threshold of 3000. We gated the bacterial population as those events with an FL1-A > 10^4^ and FSC-A < 10^5^. The raw flow cytometry data files are available at https://doi.org/10.6084/m9.figshare.15104409.

### Alignment and phylogentic tree

The selection-switch dataset^12^ was acquired directly from the author, in which the innoculum samples were discarded. The OTU reference sequences were aligned with SINA version 1.6.1^44^ using the SILVA Release 138 NR 99 SSU dataset^45^. Using this aligment, the phylogentic tree was constructed by neighbour-joining using MEGA X^46^ with default parameters.

### Filtering and preprocessing

The read counts for each OTU in each sample were divided by the total number of reads for the sample, generating relative abundances. Thereafter, all OTUs having a maximum abundance (across all samples) below a certain threshold, were removed. Three levels of filtering thresholds were applied: High level at 5 · 10^−3^, medium level at 1 · 10^−3^ and low level at 5 · 10^−4^. For obtaining estimates of absolute abundances, the relative abundances were scaled by the estimate of total bacterial cell count for each sample. The `phyloseq` package^47^ and the `R` programming language^48^ facilitated this procedure. In addition, we wrote an R-package named `micInt` (available at https://github.com/AlmaasLab/micInt) to facilitate and pipeline the analysis.

### Similarity measures and addition of noise

In total seven similarity measures were used for assessing co-occurrence between the OTUs. These were: Pearson correlation, Spearman correlation, Squared Euclidean similarity, Jaccard similarity, generalized Jaccard index, cosine similarity and Bray-Curtis similarity. Of theses similarity measure, only the Pearson correlation and Spearman correlation are considered in this article. A similarity measure, as referred to in this article, can be thought of as a function 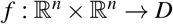, where *D* = [0,1] for the unsigned similarity measures (Squared Euclidean, Jaccard, cosine, and Bray-Curtis) and *D* = [−1,1] for the signed similarity measures (Pearson and Spearman). In this regard, *f* (**x**, **y**) is the similarity of two abundance vectors **x** and **y** belonging to different OTUs. Given two vectors **x** and **y** of abundances, normally distributed noise was added to each of the abundance vectors and the similarity measure has invoked thereafter. This is, given a similarity measure *f*, the similarity between the abundance vectors after adding noise is given by:

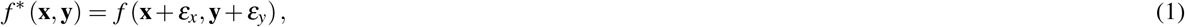

where *ε_x_* and *ε_y_* are compontentwise independent and normally distributed with mean zero and variance *γ*^2^. The level of noise γ was determined by the smallest non-zero relative abundance *x*_min_ in the dataset and a fixed constant *s* called the *magnitude factor*, such that *γ* = *s* · *x*_min_. For no noise, *s* = 0, for low noise *s* = 1, for middle noise *s* = 10 and for high noise *s* = 100.

### Network creation

Significance of the pairwise OTU associations were determined by the ReBoot procedure introduced by Faust *et al*.^23^ and shares the underlying algorithm used in the CoNet Cytoscape package^49^. This approach accepts a dataset of microbial abundances and a similarity measure, and evaluates for each pair of OTUs in the dataset the null hypothesis *H*_0_: “The association between the OTUs is caused by chance”. By bootstrapping over the samples, the similarity score of each pair of OTUs is estimated, forming a bootstrap distribution. By randomly permuting the pairwise abundances of OTUs and finding the pairwise similarity scores, a bootstrap distribution is formed. The bootstrap and permutation distribution are then compared with a two-sided *Z*-test (based on the normal distribution) to evaluate whether the difference is statistically significant. For this, the *z*-value, *p*-value and *q*-value (calculated by the Benjamini-Hochberg-Yekutieli procedure^50^) are provided for each pair of OTUs in the dataset. Our ReBoot approach is based on the `R`-package `ccrepe`^51^, but is integrated into the `micInt` package with the following major changes:

- The original ReBoot uses renormalization of the permuted abundances to keep the sum-to-constant constraint. Whereas this is reasonable to do with relative abundances, our modified version enables turning this feature off when we analyse data with absolute abundances.
- Optimizations have been made to memory use and CPU consumption to enable analyses of large datasets.
- In contrast to the usual ReBoot procedure, networks generated by the different similarity measures are not merged by *p*-value, but kept as they are.

For our analysis the number of bootstrap and permutation iterations was set to 1000. All OTUs being absent in more than 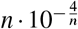 samples, where *n* is the total number of samples, were excluded through the `errthresh` argument but still kept for renormalization (if turned on). The associations were made across all samples, even the ones belonging to a different selection group or nutrient supply.

### Dynamic PCoA visualization

All samples in the dataset were used for PCoA ordination, where the Bray-Curtis distance metric between the samples was applied before creating the decomposition. After the ordination was computed, the samples were divided into four facets based on their combination of current selection regime and nutrient supply. Finally, all samples belonging to the same microcosm were connected by a line in chronological order and the line was given a separate style based on the nutrient supply and coloured to visually distinguish it from the two other replicate microcosm within the same facet.

### Network visualization

The networks were plotted by the R package `igraph`^52^. Network modules were found by the walktrap^20^ algorithm implemented in `igraph` with the setting `steps=20`, including the positive edges only. Later, the negative edges were added and the networks plotted with the community labelling.

The time dynamics of the networks were visualised by taking the former network and adjusting the node colour and size, as well as the edge colour. For this, a certain combination of selection group (i.e RK) and nutrient supply (i.e H) was chosen. Further, let *x_i,j,k_* be the abundance of OTU *k* at sampling day *i* in microcosm *j*. As there are three replicates, we have that *j* = 1,2,3. If the underlying network was created by Pearson correlation, we denote the day mean *x_i,.,k_* as the average over the replicates, this is:

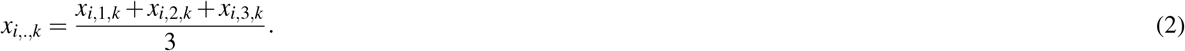

The time series mean of OTU *k*,*x...._k_* is the mean of these daily means over all sampling days,

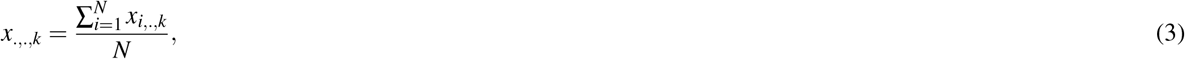

where *N* denotes the number of sampling days. Furthermore, we have the associated standard deviation *σ_k_* as given by:

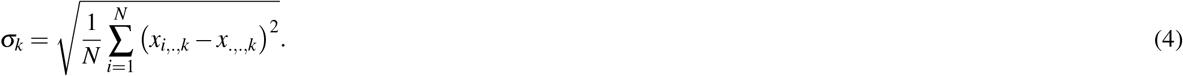

The z-value of the abundance of OTU *k* at day *i* is then:

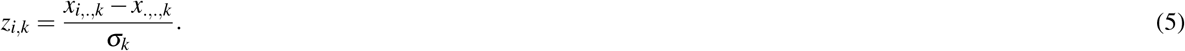

This value is used in the mapping of the node sizes and colours. The node for OTU *k* at sampling day *i* has the size *a*+*b* · |*z_i_,_k_*|, where *a* and *b* are constants. Furthermore, the same node is coloured:

- Black if *z_i,k_* < −1. This indicates that the OTU that day had a lower abundance than the average.
- Grey if −1 ≤ *z_i,k_* ≤ 1. This indicates that the OTU that day had about the same abundance as the average.
- Orange if *z_i,k_* > 1. This indicates that the OTU that day had a higher abundance than the average.

Furthermore, the edge colour are dependent on the product of the two participating nodes. Hence, the edge between OTU *k* and OTU *l* at day *i* will have the colour:

- Red if *z_i,k_ · z_i,l_* < −0.3. This shows a contribution to a negative interaction.
- Gray if −0.3 ≤ *z_i,k_ · z_i,l_* ≤ 0.3. This shows no major contribution of neither a positive nor negative interaction.
- Blue if *z_i,k_ · z_i,l_* > 0.3. This shows a contribution to a positive interaction.

Our approach is motivated by the fact that the Pearson correlation *ρ_k,l_* of the day means of OTU *k* and OTU *l* is given by:

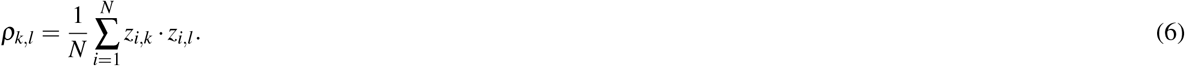

For the Spearman correlation, the visualization is based on the rank of each of the OTU abundance values in a sample. Hence, instead of using the raw abundances *x_i,j,k_* in the calculation of the day mean, the ranks *r_i,j,k_* are used instead, and all subsequent calculations and mappings are the same. In a scenario when there is only one replicate, the quantity *ρ_k,l_* would then be the Spearman correlation of the abundances of OTU *k* and OTU *l*.

## Acknowledgements

Thanks to Olav Vadstein for being helpful with explaining the experimental procedures and supply input on how the data could be interpreted and processed. J.P.P. and E.A. are grateful to ERA CoBioTech project CoolWine and Norwegian Research Council grant 283862 for funding.

## Author contributions statement

E.A. and J.P.P. conceived the project. M.S.G provided the data and comments to how to interpret it. J.P.P wrote the software, did the analysis, created visualisations and wrote the first draft of the paper. All authors contributed to and accepted the final version of the paper.

## Additional information

### Availability of source code and data

The `micInt` package used for the analysis is available at GitHub https://github.com/AlmaasLab/miclnt. The selection-switch dataset is provided as an experiment-level `phyloseq` object in a R binary dataset file in the supplementary materials. The source code and intermediate datafiles for the analysis could not be provided as supplementary material due to large size of the files, but are nevertheless available upon direct contact with the corresponding author.

